# Basal Cambrian soft-bodied segmented bilaterians preserved as microbial pseudomorphs

**DOI:** 10.1101/2024.07.03.601876

**Authors:** Xiaoguang Yang, Deng Wang, Zhiliang Zhang, Xing Wang, Jie Sun, Wenjing Hao, Yiqun Liu, Kentaro Uesugi, Tsuyoshi Komiya, Jian Han

## Abstract

Before Cambrian Stage 3, unambiguous body fossils of segmented bilaterians were rare, severely hampering our understanding of the early history of such important animals. Here we report a variety of microfossils with quintessential features such as paired appendages, dorsoventral and anteroposterior differentiations from the basal Cambrian Fortunian of South China, representing the earliest known three-dimensional body fossils of segmented bilaterians. These fossils were all microbial pseudomorphs built up by secondarily phosphatized bacteria aggregations, testifying microbial pseudomorph could serve as a novel and important pathway to preserve tiny, fragile bilaterian progenitors. This finding unveils a diversified segmented bilaterian world at the very beginning of Cambrian and would arouse a more comprehensive perspective on the early evolution of bilaterian body plans.

## Introduction

Segmented bilaterian invertebrates, i.e., panarthropods and annelids, have been one of the most successful and widespread animal groups on Earth since the Cambrian Period, about half billion years ago. Their origin and early evolution are hotly discussed topics in palaeontologic research (Buatois *et al*. 2014; Chen *et al*. 2018; Chen *et al*. 2019; Peterson *et al*. 2008; Steiner *et al*. 2004). The earliest records of trace fossils and body fossils of segmented bilaterians could probably date back to Late Ediacaran (Chen *et al*. 2018; Chen *et al*. 2019; Glaessner 1958) and molecular clock data suggested a further deeper origin (Erwin 2020; Erwin *et al*. 2011). However, between these late Proterozoic cases and the great radiation of panarthropods (Budd 1998; Caron & Aria 2017; Ortega-Hernández *et al*. 2017; Ou & Mayer 2018; Vannier & Martin 2017;Yang *et al*. 2015; Zhang *et al*. 2016) and annelids (Chen *et al*. 2020; Han *et al*. 2019; Nanglu & Caron 2021; Parry & Caron 2019) onwards Cambrian Stage 3, there is a notable absence of segmented bilaterian fossil records, though trace fossils implied bilaterian with paired appendages did exist at least during the Ediacaran–Cambrian transition (Buatois *et al*. 2014). Moreover, among the early Cambrian exceptionally preserved fossil assemblages that yielded many phosphatized soft-bodied animals (Steiner *et al*. 2004; Bengtson & Yue 1997; Han *et al*. 2013; Han *et al*. 2017; Liu *et al*. 2014), body fossils of uncontroversial segmented bilaterians remain unknown to date. Probably due to the soft and microscopic nature of primitive forms of segmented bilaterians during this period (Anderson *et al*. 2023; Budd & Jackson 2016), they were difficult to be preserved in ordinary way. On the other hand, microbial colonies could replicate the gross morphologies (forming microbial pseudomorphs) of the original soft tissues, usually muscle of the host macro-animals, after consuming them (Allison 1988; Briggs 2003; Wilby & Briggs 1997). In the setting of phosphatized microfossils, microbial pseudomorphing had been regarded as an effective pathway to preserve embryos (Eagan *et al*. 2017; Raff *et al*. 2014; Raff *et al*. 2008; Raff *et al*. 2013), but no relevant body fossil of micro animal was reported yet.

Here we present a dozen of earliest-known segmented bilaterian body fossils (Figures 1 and 2 and figure supplements 2 to 4 and supplementary movies 1-7) from the Kuanchuanpu Formation of South China, belonging to the basal Cambrian Fortunian (refer to Materials and Methods) and all of them were preserved as three-dimensional microbial pseudomorphs. These segmented bilaterian fossils give us the crucial missing piece of the evolutionary puzzle and validate that such animals had well developed and highly diversified at the basal Cambrian for the first time. While in terms of taphonomy, it confirms the potential of microbial pseudomorphing to preserve micro animals (Eagan *et al*. 2017; Raff *et al*. 2013; Butler *et al*. 2015) and further explains how this mechanism helped such small and soft bilaterian ancestors overcome the adverse preserving conditions and became a unique window for exceptional preservation soft-bodied micro-animals.

**Figure 1.**
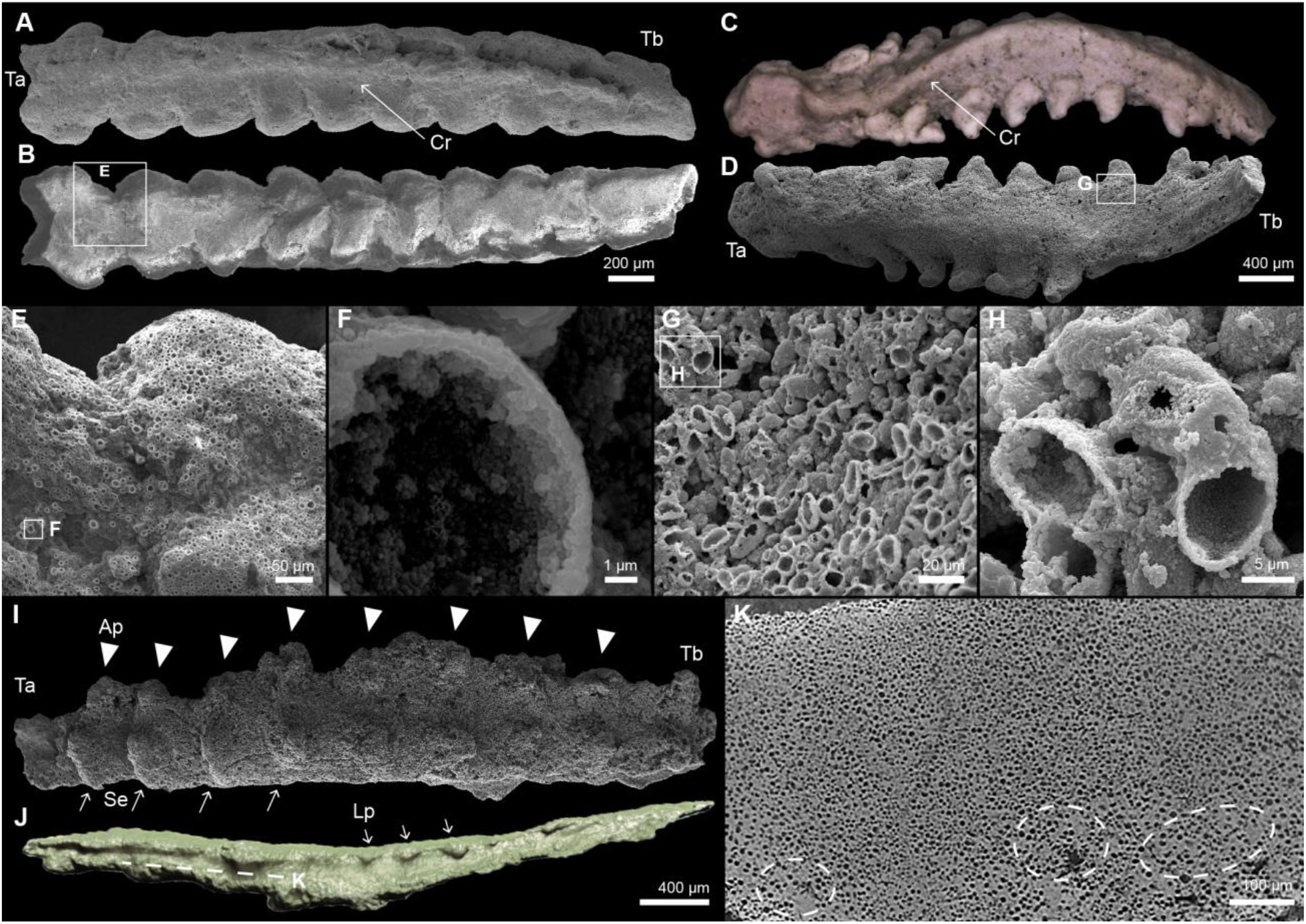
Appendage-bearing bilaterians preserved as microbial pseudomorphs from the Cambrian Kuanchuanpu Formation, South China. (**A**) Dorsal view of ELIXX92-17, with a blunt terminus-a (Ta), a slightly tapered terminus-b (Tb) and a central ridge (Cr) along anterior-posterior axis. (**B**) Ventral view of ELIXX92-17, showing nine pairs of appendages. Boxed area enlarged in (E). (**C**) Dorsal view of ELIXX88-1, with a blunt terminus-A (Ta), a gradually tapered terminus-b (Tb) and a central ridge (Cr) through its body. (**D**) Ventral view of ELIXX88-1, showing nine pairs of appendages. Boxed area enlarged in (G). (**E**) Compact aggregation of spherical micrograins. Boxed area enlarged in (F). (**F**) Detailed structure of a spherical micrograin as a thin-walled vesicle built by phosphate nanocrystals. (**G**) Compact aggregation of ellipsoidal micrograins. Boxed area enlarged in (H). (**H**) Detailed structure of an ellipsoidal micrograin, similar with the spherical counterparts. (**I**) Ventral view of ELIXX96-21, showing 8 pairs of short cylindrical appendages (Ap) and 4 recognizable segments (Se). Within each segment, the biased location of an appendage pair to terminus-a (Ta) suggest an anteroposterior differentiation. (**J**) Three-dimensional model of ELIXX96-21 generated by micro-CT data, showing a flatten body with lateral protrusions (Lp). (**K**) Tomographic view of area marked by dotted line in (J), showing compact contacts between microgains or solid spacing areas (dotted line circles).

**Figure 2.**
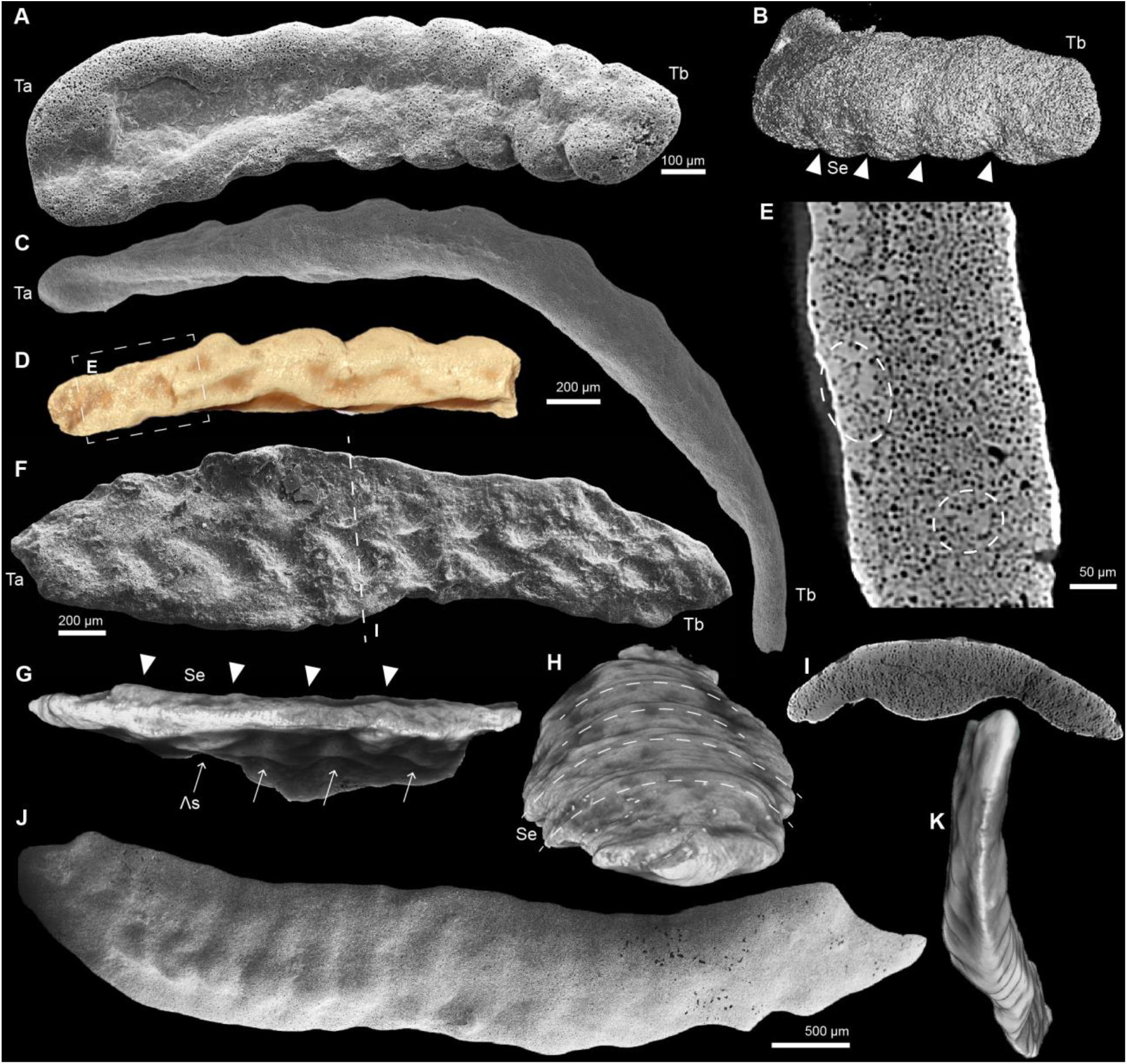
Segmented bilaterian pseudomorphs from the Cambrian Kuanchuanpu Formation, South China. (**A**) Ventral view of ELIXX59-71, showing a blunter terminus-a (Ta) and thinner terminus-b (Tb), with a distinct groove along body axis and five pairs of recognizable stubby appendages. (**B**) Three-dimensional CT model with dorsal view of ELIXX59-71, showing clear segments (Se) correlating with each appendage pair. (**C**) Dorsal view of ELIXX56-367, with a rounded terminus-a (Ta) and a long gradually tapered terminus-b (Tb). (**D**) Three-dimensional CT model with ventral view of cracked part of ELIXX56-367, showing a series of bulges arranged along the body axis. (**E**) Tomographic view of area marked by dotted line box in (D), showing homogeneous aggregation of spherical micrograins and solid cements between micrograins (dotted line circles). (**F**) Ventral view of ELIXX73-713 with a wider terminus-a (Ta) and a narrower terminus-b (Tb), presenting pairs of “Λ” shaped structures (Λs), possibly related to appendages. (**G**) Three-dimensional CT model of left part of ELIXX73-713 with lateral-ventral view, indicating each “Λ” structure (Λs) correlated with the segments (Se) on the back. (**H**) Dorsal view of the three-dimensional CT model in (G), showing segments (Se) marked by transverse ridges. (**I**) Tomographic view of section marked by dotted line in (F), demonstrating a bilateral body. (**J**) Lateral view of ELIXX84-262, with a segmented trunk. (**K**) Three-dimensional CT model of ELIXX84-262, showing the bilateral symmetry of body segments.

## Results

The specimens described here all show up as elongated bodies with disparate shapes and substructures. They are millimeters in size, exhibiting a clear bilaterial symmetry (Figures. 1 and 2 and figure supplement 4) and distinguishable differentiations in two terminations along the elongated axes. Occurrences of ridges (Figure 1A and C), grooves (Figure 2A and figure supplements 4A, C, E, G, I) and paired protuberances which are installed on or tilt towards one side (Figures 1 and 2A and F and figure supplements. 3A to C and I and 4) indicate two differentiated sides for their bodies. In addition, most of the specimens show flexure in varied degrees (Figures. 1A to D and J and 2 and figure supplements 3A and 4), suggesting they were pliable before fossilization. Considering above features, these specimens stand out from the ubiquitous skeletal fragments in the Kuanchuanpu fossil assemblage and could be comparable to soft-bodied bilaterian animals with anteroposterior and dorsoventral differentiations, and multiple pairs of appendages. Furthermore, tangible segmentation (Figures. 1I and 2B, G, H and J) or paired appendage arrangements (Figures. 1A to D, I and 2A and figure supplement 4) imply most of them had segmented trunks.

When zoomed in, all specimens take on the form of compact aggregations of micron-scale granules. Each grain maintained a clear boundary without interpenetration or inclusion. The granules contacted with each other tangentially (Figures. 1E, G and K and 2, E and I) or with spacings filled by infarctate cements (Figures. 1K and 2E and I). This constitution pattern was also valid to the interior so that the original anatomies of animals were all transformed into solid, homogeneous bulks (Figures. 1K and 2E and I).

The constitutive micrograins of these specimens generally appeared to be thin-walled, consisting of phosphate nanocrystals. We identify two morphotypes of them: Type-A: small spherical grains (diameter 7–12 μm, average ∼9 μm) (Figure 1E and F and figure supplement 2B, F and H and S4) and Type-B: small ellipsoidal grains (averaging long axis 14 μm, short axis 7 μm) (Figure 1G and H and figure supplements 2D, E, G and H and 4H). Most specimens are comprised of single type of granule, while minor mixture of type B within the masses dominated by Type-A occasionally occurred in two specimens (figure supplement 3D and 4H).

## Discussion

### Biogenic microstructures and pseudomorphing mechanism

The biogenic origin of constitutive grains of specimens described here was convincing because of their regular shapes, stable dimensions, and no sign of intersection and most important, the distinctive resemblances between their aggregations and varied animals (Figures 1, 2 and figure supplement 2 and table supplement 1). By contrast, under phosphatization settings, biomorphic microparticles generated by diagenesis processes exhibited wide and random distribution of size, needle-like radiating crystallites, mutually intersected framboids, and formation of irregular or scattered aggregations (Baturin & Titov 2006; Cosmidis *et al*. 2013; Schiffbauer *et al*. 2012). Therefore, these micrograins could be regarded as fossilized microorganisms and their aggregations are microbial pseudomorphs of the host animals.

The exquisite preservation of these micrograins as hollow envelopes further reinforced their microbial affinity. Such preservation is the trait of the mineralized Gram-negative bacteria and can be a signature of microbial fossils (Cosmidis *et al*. 2013). The anionic organic polymers within the periplasm of Gram-negative bacteria cell walls provide preferential nucleation sites for Ca-phosphate, so their fossilization started with the mineralization of the cell walls (Benzerara *et al*. 2004; Fortin *et al*. 1997; Southam & Donald 1999) and subsequently left encrustations perfectly retained the shapes and dimensions of the original microbes (Cosmidis *et al*. 2013). This scenario coincided with the fine particle texture on the surfaces and tangent contacts of the Kuanchuanpu micrograins. (Figures. 1E to H and 3A to C). Meanwhile, modern taphonomic experiments showed that proteobacteria, especially gamma proteobacteria, which also belonged to Gram-negative strains, were the major pseudomorph makers (Eagan *et al*. 2017; Raff *et al*. 2014; Raff *et al*. 2008; Raff *et al*. 2013). And the solid cements between some grains (Figures. 1K and 2E and I) were interpreted as Ca-phosphates precipitation possibly induced by the extracellular polymeric substances within microbial colonies (Raff *et al*. 2008).

Although both endogenous bacteria living in the digestive tracts of host animals and exogenous bacteria from environments could form pseudomorphs (Eagan *et al*. 2017; Raff *et al*. 2013; Butler *et al*. 2015), we inferred that the microbes responsible for the studied fossils were as exogenous, based on the single-strain dominated microbiota and the absence of internal structures. Because endogenous bacteria flora was complex, containing varied morphotypes, and usually led to preservation of anatomical structures of the host animals, such as guts (Eagan *et al*. 2017; Raff *et al*. 2013; Butler *et al*. 2015).

Three steps were proposed to interpret the forming process of Kuanchuanpu pseudomorph. Firstly, exogenous heterotrophic Gram-negative bacteria, probably proteobacteria invaded an animal carcass from body openings (mouth, anus) (Figure 3D). Next, they rapidly proliferated within the body cavity, consuming the internal tissues and outcompeting the endogenous microbial flora, just as typical exogenous pseudomorph maker *Pseudoalteromonas tunicata* (gamma proteobacteria) did (Eagan *et al*. 2017). The dense, single-strain dominated microbial colonies were then crammed against the more tenacious integument (possible cuticle) and replicated animal profile (Figure 3E and figure supplement 5). Finally, the formed pseudomorph, with extraordinary stability (Raff *et al*. 2013), persisted through further degradation which might lead to failure of the integument and mineralized as fossils (Figure 3F). In our cases, Type-A microbes were more active species, with the domination in both mixed microbiotas (figure supplement 3E and 4H) and total specimen amount, implying competition also occurred among able pseudomorph makers, which testified the complicated interaction between bacteria strains observed in taphonomy experiments (Eagan *et al*. 2017; Raff *et al*. 2013).

**Figure 3.**
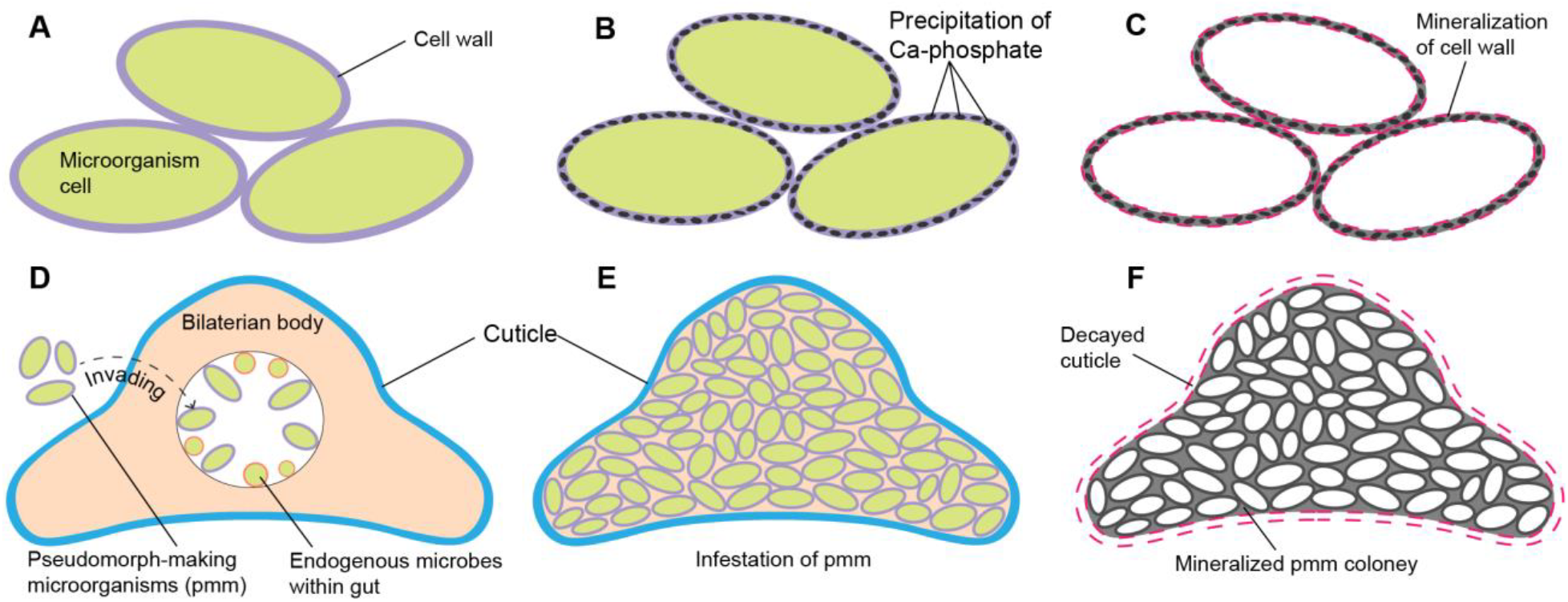
Mechanism of forming an animal body pseudomorph. (**A** to **C**) The mineralization of pseudomorph-making microbe cells started within their cell walls and preserved as hollowed envelope-like fossils. (**D** to **F**) The interaction of endogenous and exogenous microbiotas and the pseudomorphing and fossilizing progress of a bilaterian body.

Concluded from the microbial affinity and pseudomorphing mechanism, three major advantages can explain the superiority of pseudomorphing over direct phosphatization of soft-bodied animals themselves: i) The more feasible templates for mineralization provided by cell wall structures of specific bacteria; ii) catalyzed precipitation of Ca-phosphates induced by both bacteria cells (Cosmidis *et al*. 2013) and extracellular matrix (Raff *et al*. 2008); iii) the extraordinary stability of formed pseudomorphs to prevent other bacteria which rapidly destroy ordinary soft tissues and keep the pseudomorphs long enough for later mineralization (Raff *et al*. 2013).

### Implications on early evolution of segmented bilaterians

The pseudomorphing preservation could be regarded as kind of “internal mold”, so the fidelity of the gross body plan preservation was secured, although internal anatomies, integumental textures and subassemblies (sclerites, spines, setae) were usually lost, making accurate phylogenetic assessments difficult. The remaining torsos demonstrated quite a morphological disparity and might suggest different locomotion styles: i) segmented bilaterians with pairs of appendages (Figures. 1A, C and I and 2A and F and figure supplement 4A, C, E and I). The appendages showed no specialization, but their positions and orientations varied among representative specimens. Those grew vertically to animals ventral side were stubby and stout (Figures. 1I and 2A and F and 4B and figure supplement 4A, C and G) while longer limbs extended laterally or semi-laterally (Figures. 1A to D and 4E and H and figure supplement 3A, B and I), suggesting these micro-benthos could not lift body up effectively and probably lived on crawling over the sea floor (Figure 4J), thus cross-validating the presence of earliest Cambrian appendage-bearing bilaterians previously marked only by trails on the biomat grounds (Buatois *et al*. 2014); ii) segmented bilaterian without appendage (Figure 2J and K). This fossil had laterally flattened body with longitudinally arranged segments. The recognizable segments appeared to be homonomous but with different shapes in two ends, together with the body cross section (figure supplement 3K) implied a dorsoventral differentiation; iii) elongate vermiform body with a blunter terminus and a gradually tapered end, indicating an anterior-posterior axis. A series of bulges arranged along the body axis and showed vaguely bilaterial symmetry and segments (Figure 2C and D).

**Figure 4.**
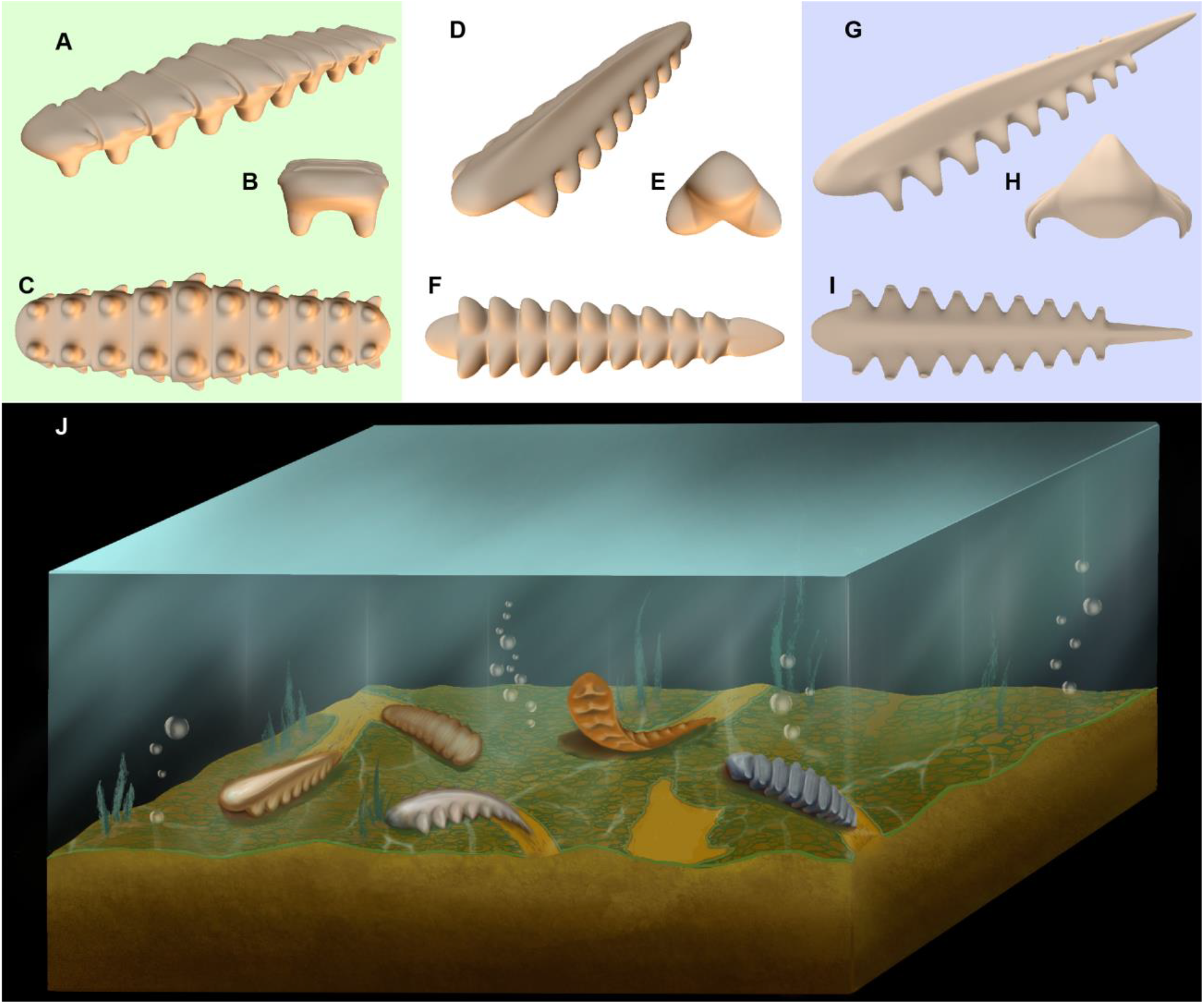
Diversity of early Cambrian bilaterian animals in morphology and ecology. (**A** to **I**) Three-dimensional reconstruction model of ELIXX96-21, ELIXX92-17 and ELIXX88-1, with overall view (A, D and G), front view (B, E and H) and ventral view (C, F and I). Their appendages extended to the ventral side vertically in the (B), semi-laterally in the (E) and laterally in the (H). (**J**) Presumed palaeoecological reconstruction art with the typical early Cambrian bilaterian animals. From left to right: ELIXX92-17, ELIXX59-71, ELIXX88-1, ELIXX73-713, ELIXX96-21.

With the remarkable bilaterality, body segments and paired appendages, most of these fossil animals could be unequivocally assigned to panarthropods or annelids, thus extending their occurrence from Cambrian Stage 3 into the dawn of the Phanerozoic, approximately 20 million years forward. The soft and microscopic bodies with pliable but somehow tenacious integument might present the expected progenitors of segmented bilaterians, which were thought to be soft and small (Anderson *et al*. 2023; Budd & Jackson 2016) and long missing during the early Cambrian. Thanks to this new exceptional preservation pathway as microbial pseudomorphing, the taphonomy bias (dos Reis *et al*. 2015) could be overcame to a certain extent, which fills the cryptical gap of body fossils interval of segmented bilaterian (Budd & Jackson 2016) between Late Ediacaran (Chen *et al*. 2019; Glaessner 1958) and Cambrian Stage 3 (Budd 1998; Caron & Aria 2017; Ou & Mayer 2018; Vannier & Martin 2017;Yang *et al*. 2015; Zhang *et al*. 2016; Chen *et al*. 2020; Han *et al*. 2019; Nanglu & Caron 2021; Parry & Caron 2019), suggesting the Cambrian Explosion might root deeper than we expected. The rudimentary traits of these animals, such as homonomous body segments, non-specialized appendages, and limited motility suggests they were still in early evolutionary stage of segmented bilaterian, while their considerable diversity in morphology implies much earlier divergences than Cambrian and possibly illuminate the subsequent great radiation of bilateral animals. In conclusion, the new bilaterians discovered here represented key transitional groups in the critical geological transition as Cambrian Explosion, mending a major imperfection for the fossil record of early animal evolution and contributing to bridge the gap between molecule clock hypothesis and geological data (Erwin 2020; Erwin *et al*. 2011; Budd & Jackson 2016; Budd & Mann 2020).

## Materials and Methods

### Geological setting

The Xixiang area was palaeogeographically located on the northwestern margin of the Yangtze Platform during the Ediacaran and Cambrian periods (figure supplement 1A). The Kuanchuanpu Formation is well exposed at this section, with a total thickness of ca. 21 m. It disconformably underlies the black shale of the Lower Cambrian Guojiaba Formation and overlies the Neoproterozoic Upper Dolomite Member of the Dengying Formation in a para-unconformity. From lower to upper, the Kuanchuanpu Formation can be subdivided into 4 Beds as follows: (1) grey blocks fmicrosparitic limestone (0.8 m); (2) grey and white blocks of phosphorus limestone (2.2 m); (3) thick-bedded dark grey microsparitic limestone (17.4 m); (4) thin-bedded light grey silty-fine dolomitic limestone (0.6 m) (figure supplement 1B).

The sampling horizon is Bed 2. The fossil assemblage includes *Protohertzina anabarica, Canopoconus calvatus, Conotheca subcurvata, Olivooides multisulcatus*, and *Olivooides mirabilis*, which could be correlated with the first microfossil assemblage zone of Fortunian Stage (*Anabarites trisulcatus-Protohertzina anabarica* assemblage zone). The estimated age of sampling horizon is approximately 535 Ma (figure supplement 1B) (Yang *et al*. 2017).

### Materials

All specimens were recovered from the Bed 2 of phosphorus limestone of the Kuanchuanpu Formation at the Zhangjiagou Section, Dahe Town, Xixiang County, Hanzhong City, Shaanxi Province, China (fig. S1). Rock samples were treated with 8–10% acetic acid solution and phosphatized fossils were then selected from the residues using a binocular microscope.

### Scanning electron microscopy (SEM)

Selected fossils were coated with gold then observed with an FEI Quanta 450 FEG Scanning Electron Microscope (SE Mode, high vacuum, 20 kV, spot size 4.0) at State Key Laboratory of Continental Dynamics, Northwest University, Xi’an, China.

### X-ray computed microtomography and 3D reconstruction

High-potential specimens were analyzed with a ZEISS Xradia-520 micro-computed tomography (50 kV/4W, 80 μA, 1.10×1.10×1.10 to 4.23×4.23×4.23 μm^3^ pixel^-1^ depended on fossil size) at the State Key Laboratory of Continental Dynamics (SKLCD), Northwest University, China. One specimen ELIXX59-71 was analyzed at SPring-8 in Hyogo, Japan (23 KeV, 0.49×0.49×0.49 μm^3^ pixel^-1^). X-ray microtomography data were processed through ORS Dragonfly Pro (v. 2022.1.0.1259) software for detailed tomographic analysis and making three-dimensional visualization pictures and videos.

### Measurements

Statistical analyses were based on the micrograins which exhibit relatively intact three-dimensional outlines and diameters could be determined. Measurements for the grain size were performed by ImageJ 1.53K with selected SEM images.

## Acknowledgments

We thank J. Luo (NWU) for assistance with fossil collection, X. Liu (NWU) and W. Zhou (XAUT) for the artistic reconstructions, Profs. J. Paterson (University of New England) and Q. Ou (China University of Geosciences Beijing) for constructive comments during draft writing and editing.

## Author contributions

Conceptualization: XY, JH

Field work: XY, XW, JH

Micro CT analyses and visualization: WH, JS

SPring-8 analyses and data reconstruction: KU, TK

Writing – original draft: XY, DW, ZZ

Writing – review & editing: XY, JH, DW, ZZ, XW, YL

## Data and materials availability

Data of Micro CT and SPring-8 analyses are available at figshare (https://figshare.com/s/b1a9aa032d009fdbeaac). All figured specimens are housed in the collections of the Early Life Institute of Northwest University and can be accessed by contacting Jian Han. All data are available in the main text or the supplementary materials.

## References

Allison PA (1988) Phosphatized soft-bodied squids from the Jurassic Oxford Clay Lethaia 21: 403–410 doi: 10.1111/j.1502-3931.1988.tb01769.x

Anderson RP, Woltz CR, Tosca NJ, Porter SM, Briggs DEG (2023) Fossilisation processes and our reading of animal antiquity Trends In Ecology & Evolution 38: 1060–1071 doi: 10.1016/j.tree.2023.05.014

Baturin GN, Titov AT (2006) Biomorphic formations in recent phosphorites Oceanology 46: 711– 715 doi:10.1134/S0001437006050110

Bengtson S, Yue Z (1997) Fossilized metazoan embryos from the earliest Cambrian Science 277: 1645–1648 doi: 10.1126/science.277.5332.1645

Benzerara K, et al. (2004) Biologically controlled precipitation of calcium phosphate by Ramlibacter tataouinensis Earth And Planetary Science Letters 228: 439–449 doi: 10.1016/j.epsl.2004.09.030

Briggs DEG (2003) The role of decay and mineralization in the preservation of soft-bodied fossils Annual Review of Earth and Planetary Sciences 31: 275–301 doi:10.1146/annurev.earth.31.100901.144746

Buatois LA, Narbonne GM, Mángano MG, Carmona NB, Myrow P (2014) Ediacaran matground ecology persisted into the earliest Cambrian Nature Communications 5: 3544 doi: 10.1038/ncomms4544

Budd GE (1998) The morphology and phylogenetic significance of Kerygmachela kierkegaardi Budd (Buen Formation, Lower Cambrian, N Greenland) Earth and Environmental Science Transactions of The Royal Society of Edinburgh 89: 249–290 doi: 10.1017/S0263593300002418

Budd GE, Jackson IS (2016) Ecological innovations in the Cambrian and the origins of the crown group phyla Philosophical Transactions of the Royal Society B-Biological Sciences 371: 20150287 doi: 10.1098/rstb.2015.0287

Budd GE, Mann RP (2020) Survival and selection biases in early animal evolution and a source of systematic overestimation in molecular clocks Interface Focus 10: 20190110 doi: 10.1098/rsfs.2019.0110

Butler AD, Cunningham JA, Budd GE, Donoghue PCJ (2015) Experimental taphonomy of Artemia reveals the role of endogenous microbes in mediating decay and fossilization Proceedings of the Royal Society B-Biological Sciences 282: 20150476 doi: 10.1098/rspb.2015.0476

Caron JB, Aria C (2017) Cambrian suspension-feeding lobopodians and the early radiation of panarthropods BMC Evolutionary Biology 17: 1–14 doi: 10.1186/s12862-016-0858-y

Chen H, et al. (2020) A Cambrian crown annelid reconciles phylogenomics and the fossil record Nature 583: 249–252 doi: 10.1038/s41586-020-2384-8

Chen Z, Chen X, Zhou C, Yuan X, Xiao S (2018) Late Ediacaran trackways produced by bilaterian animals with paired appendages Science Advances 4: eaao6691 doi:10.1126/sciadv.aao6691

Chen Z, Zhou C, Yuan X, Xiao S (2019) Death march of a segmented and trilobate bilaterian elucidates early animal evolution Nature 573: 412–415 doi: 10.1038/s41586-019-1522-7

Cosmidis J, et al. (2013) Nanometer-scale characterization of exceptionally preserved bacterial fossils in Paleocene phosphorites from Ouled Abdoun (Morocco) Geobiology 11: 139–153 doi: 10.1111/gbi.12022

dos Reis M, et al. (2015) Uncertainty in the Timing of Origin of Animals and the Limits of Precision in Molecular Timescales Current Biology 25: 2939–2950 doi: 10.1016/j.cub.2015.09.066

Eagan J, et al. (2017) Identification and modes of action of endogenous bacteria in taphonomy of embryos and larvae Palaios 32: 206–217 doi:10.2110/palo.2016.071

Erwin DH (2020) The origin of animal body plans: a view from fossil evidence and the regulatory genome Development 147: dev182899 doi: 10.1242/dev.182899

Erwin DH, et al. (2011) The Cambrian conundrum: early divergence and later ecological success in the early history of animals Science 334: 1091–1097 doi: 10.1126/science.1206375

Fortin D, Ferris FG, Beveridge TJ (1997) Surface-mediated mineral development by bacteria Reviews In Mineralogy & Geochemistry 35: 161–180

Glaessner MF (1958) New fossils from the base of the Cambrian in South Australia Transactions of the Royal Society of South Australia 81: 185–188

Han J, Conway Morris S, Hoyal Cuthill JF, Shu D (2019) Sclerite-bearing annelids from the lower Cambrian of South China Scientific Reports 9: 4955 doi: 10.1038/s41598-019-40841-x

Han J, et al. (2013) Early Cambrian pentamerous cubozoan embryos from South China PloS one 8: e70741 doi: 10.1371/journal.pone.0070741

Han J, et al. (2017) Meiofaunal deuterostomes from the basal Cambrian of Shaanxi (China) Nature 542: 228–231 doi: 10.1038/nature21072

Liu Y, et al. (2014) The oldest known priapulid-like scalidophoran animal and its implications for the early evolution of cycloneuralians and ecdysozoans Evolution & Development 16: 155–165 doi: 10.1111/ede.12076

Nanglu K, Caron JB (2021) Symbiosis in the Cambrian: enteropneust tubes from the Burgess Shale co-inhabited by commensal polychaetes Proceedings of the Royal Society B-Biological Sciences 288: 20210061 doi: 10.1098/rspb.2021.0061

Ortega-Hernández J, Janssen R, Budd GE (2017) Origin and evolution of the panarthropod head–a palaeobiological and developmental perspective Arthropod Structure & Development 46: 354–379 doi: 10.1016/j.asd.2016.10.011

Ou Q, Mayer G (2018) A Cambrian unarmoured lobopodian,† Lenisambulatrix humboldti gen. et sp. nov., compared with new material of †Diania cactiformis Scientific Reports 8: 13667 doi: 10.1038/s41598-018-31499-y

Parry L, Caron JB (2019) Canadia spinosa and the early evolution of the annelid nervous system Science Advances 5: eaax5858 doi: 10.1126/sciadv.aax5858

Peterson KJ, Cotton JA, Gehling JG, Pisani D (2008) The Ediacaran emergence of bilaterians: congruence between the genetic and the geological fossil records Philosophical Transactions of the Royal Society B-Biological Sciences 363: 1435–1443 doi: 10.1098/rstb.2007.2233

Raff EC, et al. (2008) Embryo fossilization is a biological process mediated by microbial biofilms Proceedings of the National Academy of Sciences of the United States of America 105: 19360–5 doi:10.1073/pnas.0810106105

Raff EC, et al. (2013) Contingent interactions among biofilm-forming bacteria determine preservation or decay in the first steps toward fossilization of marine embryos Evolution & Development 15: 243–256 doi: 10.1111/ede.12028

Raff RA, et al. (2014) Microbial ecology and biofilms in the taphonomy of soft tissues Palaios 29: 560–569 doi:10.2110/palo.2014.043

Schiffbauer JD, Xiao S, Sharma KS, Wang G (2012) The origin of intracellular structures in Ediacaran metazoan embryos Geology 40: 223–226 doi: 10.1130/G32546.1

Southam G, Donald R (1999) A structural comparison of bacterial microfossils vs. ‘nanobacteria’ and nanofossils Earth-Science Reviews 48: 251–264 doi: 10.1016/S0012-8252(99)00057-4

Steiner M, Zhu M, Li G, Qian Y, Erdtmann BD (2004) New early Cambrian bilaterian embryos and larvae from China Geology 32: 833–836 doi: 10.1006/dbio.2002.0714

Vannier J, Martin EL (2017) Worm-lobopodian assemblages from the early Cambrian Chengjiang biota: insight into the “pre-arthropodan ecology”? Palaeogeography, Palaeoclimatology, Palaeoecology 468: 373–387 doi: 10.1016/j.palaeo.2016.12.002

Wilby PR, Briggs DEG (1997) Taxonomic trends in the resolution of detail preserved in fossil phosphatized soft tissues Geobios 30: 493–502 doi:10.1016/S0016-6995(97)80056-3

Yang J, et al. (2015) A superarmored lobopodian from the Cambrian of China and early disparity in the evolution of Onychophora Proceedings of the National Academy of Sciences of the United States of America 112: 8678–8683 doi: 10.1073/pnas.1505596112

Yang X, et al. (2017) Euendoliths versus ambient inclusion trails from Early Cambrian Kuanchuanpu Formation, South China Palaeogeography, Palaeoclimatology, Palaeoecology 476:147–157 doi: 10.1016/j.palaeo.2017.03.028

Zhang X, Smith MR, Yang J, Hou J (2016) Onychophoran-like musculature in a phosphatized Cambrian lobopodian Biology Letters 12: 20160492 doi: 10.1098/rsbl.2016.0492

